# Plasma membrane labelling efficiency, internalization and partitioning of functionalized fluorescent lipids as a function of lipid structure

**DOI:** 10.1101/2025.05.07.652622

**Authors:** Erdinc Sezgin

## Abstract

Labeling the plasma membrane for advanced imaging remains a significant challenge. For time-lapse live cell imaging, probe internalization and photobleaching are major limitations affecting most membrane-specific dyes. In fixed or permeabilized cells, many membrane probes either lose signal after fixation or fail to remain localized to the plasma membrane. Thus, improved probes are critically needed for applications in spatial biology. In this study, we systematically compared a range of custom-synthesized and commercially available lipid-based probes for their efficiency in labeling the plasma membrane in live, fixed, and permeabilized cells. We identified a superior probe, which outperformed others due to its lipid structure. This comparison provides insights into ideal lipid probes for visualizing the plasma membrane using advanced imaging techniques.

## INTRODUCTION

Labelling lipids with fluorescent molecules for microscopy has always been a challenge. The size of lipids is often comparable to the size of fluorescent tags, hence, tagging lipids with fluorescent molecules causes significant perturbations to lipid behaviour^1^. For instance, when the fluorophore is relatively hydrophobic, it might interact with the membrane, thus the behaviour of the fluorescent lipid analogue is heavily influenced by the interactions of the fluorescent tag^2^. Furthermore, fluorescent tags close to the lipid headgroup might change the geometry of the lipid and its dynamic behaviour in the membrane^3^. Therefore, fluorescently tagged lipid analogues rarely preserve the behaviour of their native counterparts^2,4^. To overcome these issues, an inert linker (such as polyethylene glycol, PEG) is inserted between the lipid headgroup and the fluorescent tag^5^. Moreover, instead of directly labelling the lipids with the fluorophores, alternative conjugation strategies are used^6–11^. The PEG linker also helps with membrane retention as it prevents flipping of the lipids due to its size. Therefore, it is commonly used in membrane-specific probes. However, for plasma membrane labelling, we still need better probes that remain in the plasma membrane for time-lapse live cell as well as fixed and permeabilized cell imaging. Although there are efforts to find the most effective probes to label the membranous structures^12^, a systematic evaluation for plasma membrane labelling is lacking. To fill this gap, in this manuscript, we evaluated several lipid structures and two different lipid labelling strategies: direct fluorescent tagging and azide/diarylcyclooctyne (DBCO) coupling. We evaluated plasma membrane labelling, internalization and lipid domain partitioning of the probes in phase separated membranes (Figure 1).

**Figure 1.**
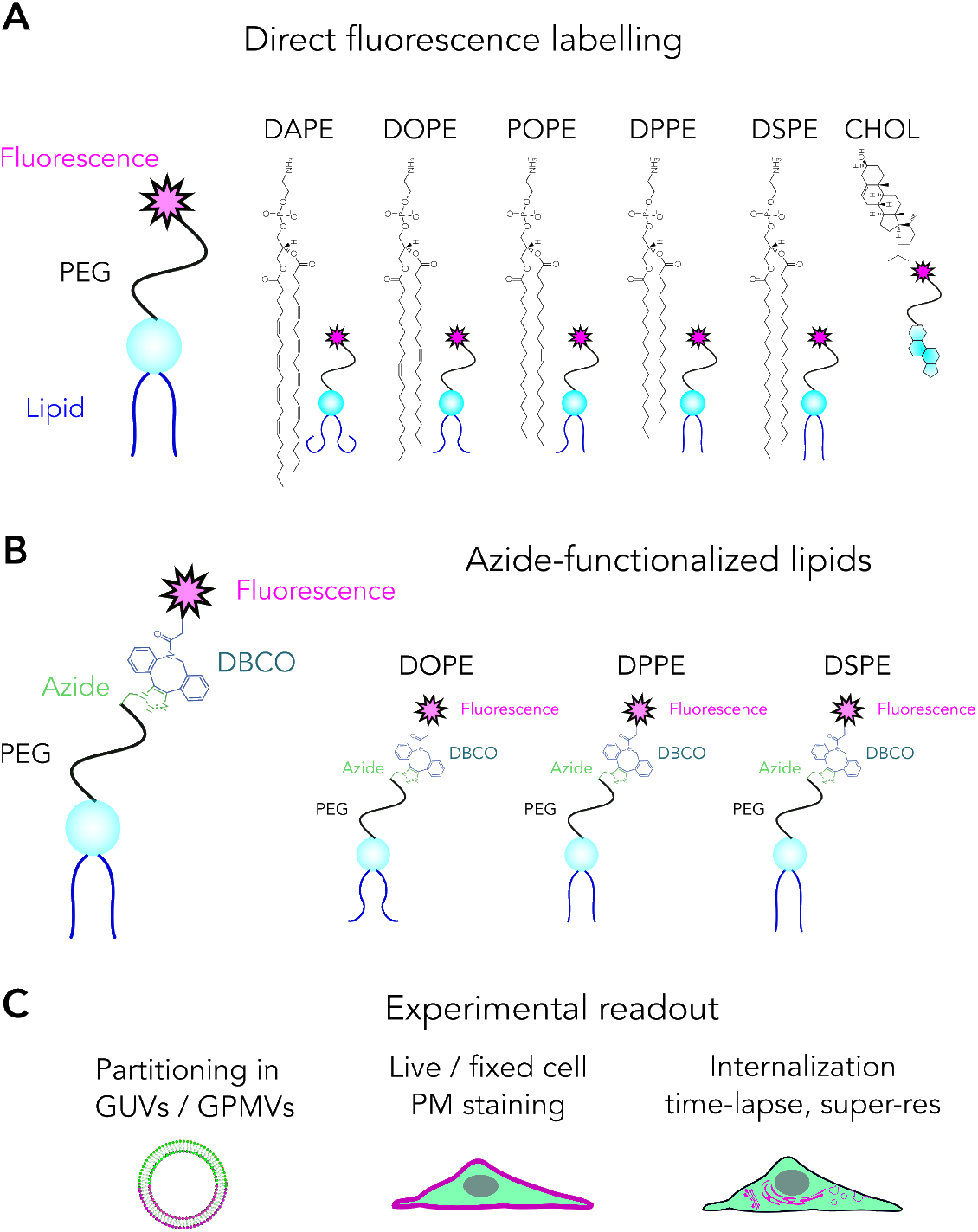
Overview of the experimental pipeline. A, B) Labelling strategies and tested lipids; and C) readouts we used in this study.

Using different labelling approaches, we identified 1,2-Dipalmitoyl-sn-glycero-3-phosphoethanolamine (DPPE) as the most suitable lipid structure for labelling the plasma membrane of two eukaryotic cell lines. We also revealed that DPPE analogues specifically partition into the saturated lipid domains better than other probes, including 1,2-distearoyl-sn-glycero-3-phosphoethanolamine (DSPE) analogues. Our data suggest that lipids with 16 carbon saturated acyl chain length is more suitable to use than 18 carbon analogues since longer acyl chain length interferes with membrane incorporation. Moreover, we found out that POPE does not partition into the liquid disordered phases as assumed previously, but rather partitioning equally in both phases. Once we identified DPPE as the most suitable lipid structure, we showed that DPPE analogues can successfully be used for long term time-lapse, fixed and permeabilized cells and for super-resolution imaging.

## MATERIALS AND METHODS

### Lipids, lipid analogs and fluorescent reporters

We purchased all native lipids from Avanti Polar Lipids. Abberior Star Red-PEG lipids were custom made by Abberior GmbH. DPPE-PEG-Azide, DSPE-PEG-Azide and DOPE-PEG-Azide were purchased from Avanti Polar Lipids. DBCO-Alexa647 was purchased from Jena Bioscience. Cell Mask was obtained from Thermofisher.

### Preparation of Giant Unilamellar vesicles (GUVs), Cells and Giant Plasma Membrane Vesicles (GPMVs)

GUVs were prepared with electroformation method^13^. Briefly, a lipid film was formed on a platinum wire from 1 mg/ml lipid mix (DOPC:SM:Chol 2:2:1). Then, GUVs were formed in 300 mM sucrose solution at 70 ºC. 10 Hz, 2 V alternative electric current was used for electroformation.

U2OS cells were maintained in DMEM medium supplemented with 10% FBS medium and 1% L-glutamine. Jurkat cells are maintained in RPMI medium supplemented with 10% FBS medium and 1% L-glutamine. GPMVs were prepared as previously described^14^. Briefly, cells seeded out on a 60 mm petri dish (with ≈70 % confluent) were washed with GPMV buffer (150 mM NaCl, 10 mM Hepes, 2 mM CaCl2, pH 7.4) twice. 2 ml of GPMV buffer was added to the cells. 25 mM PFA and 20 mM DTT (final concentrations) were added in the GPMV buffer. The cells were incubated for 2 h at 37 °C. Then, GPMVs were collected by pipetting out the supernatant.

### Fluorescent Labelling of GUVs, GPMVs and Cells

Fluorescent lipid analogues were added directly to GUVs or GPMVs with a concentration of 50 ng/mL. Azide lipids were pre-clicked with DBCO-Alexa647 in a tube. 20 ng of DBCO-Alexa647 (dissolved in water) was mixed with 1 µg of azide-modified lipid (dissolved in DMSO) for 30 min. Afterwards, the mixture was added to GUVs or GPMVs. GPMVs were incubated with these lipids at a concentration of 250 ng/ml for 1 h. Phase markers were added to GUVs and GPMVs at a final concentration of 50 ng/mL. Labelled GUVs and GPMVs were placed in the wells of BSA-coated 8-well glass bottom Ibidi chambers.

For the cell labelling, the cells were seeded on 25 mm diameter round coverslips (#1.5) in a 6 well plate 2-3 days before the measurements. The fluorescent lipid analogues were first dissolved in DMSO or ethanol with a stock concentration of 1 mg/ml. Before the labelling, the cells seeded on glass slides were washed twice with L15 medium to remove the full media. Please note that serum in the media decreases the labelling efficiency, thus it is crucial to wash out all the media from the cells. Later the fluorescent analogues were mixed with L15 medium with 1:4000 ratio (final concentration of 250 ng/ml). The cells were incubated with this suspension for 5-10 minutes at room temperature. After that, the cells were washed with L15 twice followed by imaging in the same medium. Then, confocal microscopy was performed as described below. Labelling cells with fluorescent analogues should be optimized for every cell line by changing the concentration, labelling time and labelling temperature. For CellMask labelling, it was diluted to 1:2000 and incubated with the cells for 5-10 min at room temperature.

### Confocal microscopy

GUVs were imaged in PBS, GPMVs were imaged in GPMV buffer, cells were imaged in L15 medium. Imaging was done at room temperature (21-23 °C) for fixed cells and 37 °C for live cells. All imaging was done on glass slides with thickness of 0.17 mm. Samples were imaged with a Zeiss LSM 980 confocal microscope in BSA-coated (1 mg/ml for 1 h) 8-well Ibidi glass chambers (#1.5). Abberior Star Red-labeled analogues as well as Cell Mask Deep Red were excited with 633 nm and emission collected with 650-700 nm.

Quantification of internalization has been performed in Fiji by finding the maxima (Process, Find Maxima).

### % Lo calculation

ImageJ-Line profile was used to calculate the Lo % as described in ref^15^. A line was selected which crosses the opposite sides of the equatorial plane of the GPMVs having different phases on opposite sides. Opposite sides are chosen to eliminate the laser excitation polarization artefacts. Then, %Lo was calculated as below where *I* is the fluorescence emission intensity.

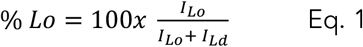

### STED imaging

STED imaging was performed using Leica FALCON system equipped with a 100x oil objective with 1.4 NA. 633 nm laser light was used to excite the probe while 650-750 nm was used to collect the emission signal. The fluorescence was depleted with 775 nm laser with 100 mW power. Images were collected with the optimal pixel number for the STED images with four-line accumulation.

## RESULTS AND DISCUSSION

### Experimental pipeline

A linker between the lipid and the fluorescent tag is essential to avoid fluorophore-induced artefacts^5^. Therefore, in all the conjugation strategies, we preserved a linker (PEG) between the lipid and the tag. Directly labelled lipids (Figure 1A) were custom-made using PEGylated lipids. These analogues were added to membranes directly. Azide-functionalized lipids (Figure 1B) were clicked with fluorescent DBCO before adding onto membranes. We used GUVs, GPMVs and live cells as target membranes to test labelling as a function of the lipid structure (Figure 1C). Using these systems, we tested partitioning of these probes in phase separated membranes, their incorporation efficiency into membranes and internalization in live cells (Figure 1C).

### Live, fixed and permeabilized cell imaging using lipids directly labelled with Abberior Star Red

We first tested the performance of different lipid probes to label of plasma membrane of cells. We hypothesized that the plasma membrane labelling depends on the choice of core lipid structure. To test this, we studied lipids with different acyl chain saturation (20:4/20:4 DAPC,18:1/18:1 DOPE, 16:0/18:1 POPE,16:0/16:0 DPPE, 18:0/18:0 DSPE and cholesterol; Figure 1A). We used a suspension and an adherent cell line to test the plasma membrane labelling efficiency of these lipids. Jurkat is an immortalized T lymphocyte cell line that grows in suspension and used for mimicking T cell signalling *in cellulo*. U-2 OS is a cancer-derived human cell lines used for various purposes such as drug screening, cell death and metabolism.

For suspension cell imaging, we stained Jurkat cells with fluorescent probes (as detailed in Methods section) for 5 minutes at room temperature and imaged them with time lapse imaging at 37 °C. All probes stained the plasma membrane of cells efficiently and remained in plasma membrane for a few hours of incubation, yet to a different extend as a function of lipid structure (Figure 2). For example, while DPPE-PEG-ASR was perfectly localized in the plasma membrane after 3h of incubation, DAPE-PEG-ASR got internalized substantially, shown by puncta structures inside the cells (Figure 2A). Internalized puncta structures suggests that internalization is driven by membrane trafficking (e.g., endocytosis). The probes also showed efficient plasma membrane labelling and retention in U-2 OS cells after hours post-labelling (Figure 2B). For both cell types, a probe without PEG linker (DOPE-ASR) internalized already after 1h post-labelling (Figure 2C) highlighting the importance of the linker in plasma membrane retention. For U-2 OS cells, we counted the number of puncta per image to quantify the internalization after 3 hours of incubation at 37 °C. The highest internalization was observed for DOPE-ASR without the PEG linker. DAPE-PEG-ASR and DSPE-PEG-ASP showed moderate; POPE-PEG-ASR and Chol-PEG-ASR showed less while DPPE-PEG-ASR showed the least internalisation (Figure 2D). This data suggests that there is lipid structure dependent internalization.

**Figure 2.**
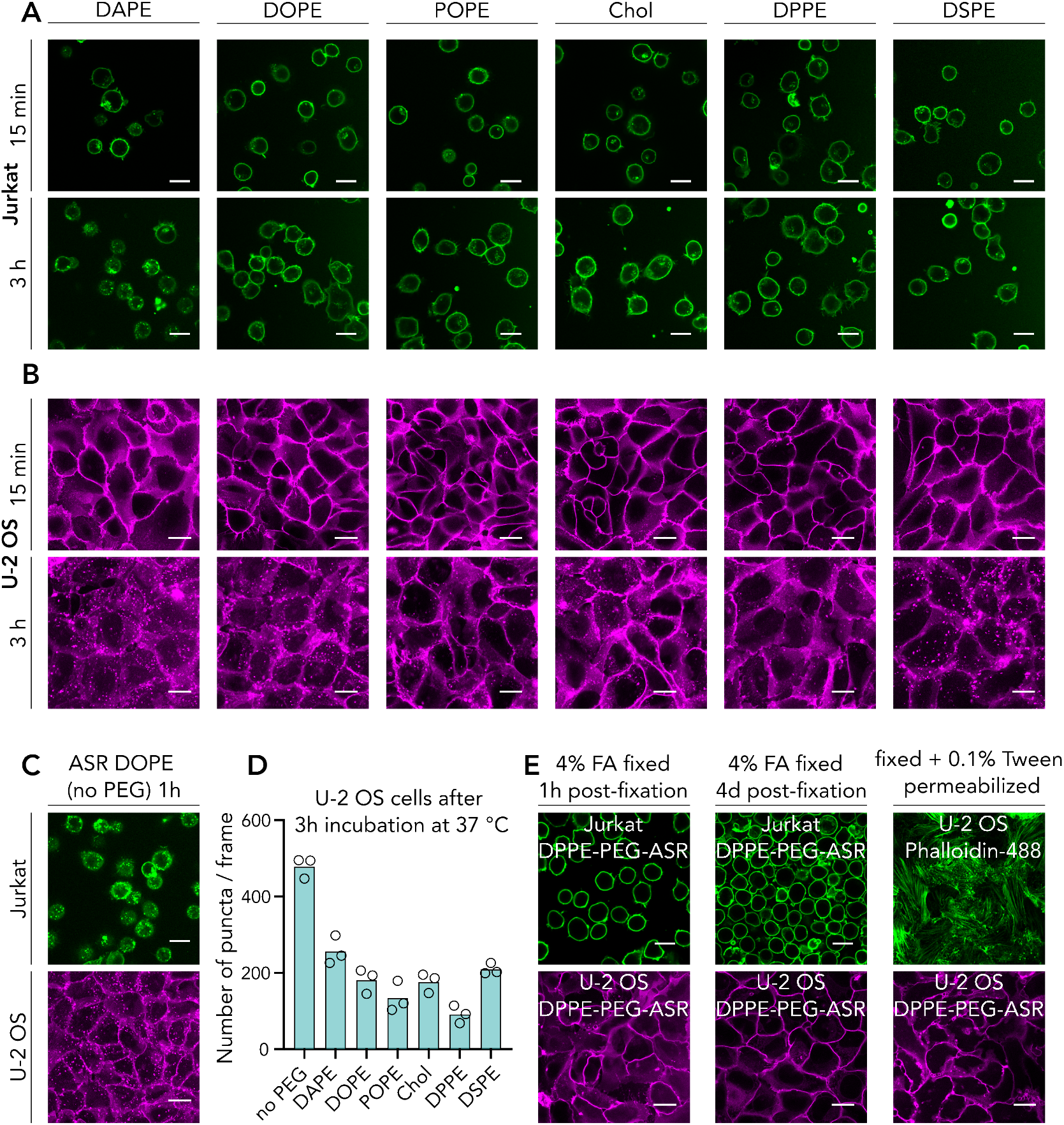
Plasma membrane staining and internalization of ASR-PEG-lipid probes in cells. A) Jurkat and B) U-2 OS live cell imaging with the probes 15 min and 3 h after the staining. C)Probe without PEG linker shows clear internalization even after 1h of incubation. D) Quantification of number of puncta per image. E) Fixed and permeabilized cells stained with DPPE-PEG-ASR. Each data point represents an image. The graph is representative of three independent experiments. Scale bars are 20 µm.

For certain experiments, cell fixation is needed. Therefore, we next investigated whether the staining can be preserved after fixation. We fixed the cells with 4% formaldehyde (FA) for 30 min at room temperature and then incubated the fixed cells with the best performing probe in live cells, DPPE-PEG-ASR. Efficient plasma membrane staining was observed after fixation (Figure 2E). The probe remained in the plasma membrane even after 4 days after labelling (Figure 2E), confirming the suitability of this probe for fixed cell imaging. Finally, certain applications require permeabilization of plasma membrane which leads to internalization of most macromolecules such as antibodies. In this scenario, it is challenging to label the plasma membrane due to detergents used for the protocol. Therefore, we tested whether these probes would sustain their plasma membrane localization after fixation with 4% FA and subsequent permeabilization with 0.5%, 0.2%, 0.1% Triton or 0.1% Tween. While Triton caused loss of probe signal at the plasma membrane (not shown), Tween-permeabilized cells sustained the membrane labelling (Figure 2E). We confirmed the permeabilization by Tween with actin labelling with phalloidin. Therefore, we conclude that these probes can be used for Tween-permeabilized but not Triton-permeabilized cells.

### Domain partitioning of lipids directly labelled with Abberior Star Red (ASR) via their head group

It was intriguing to see different internalization depending on the base lipids of the probes. Therefore, we next set out to find the underlying reason. Different endocytic routes as a function of lipid structure has been shown^16^ which can explain the varying level of internalization of different lipid analogues we studied. To test this, we evaluated the lipid domain partitioning behaviour of directly labelled lipid analogues in phase separated giant unilamellar vesicles (GUVs) and cell-derived giant plasma membrane vesicles (GPMVs). We calculated the ordered domain partitioning as liquid ordered (Lo) %, where 50% means equipartitioning; >50% means preference for ordered and <50% means preference for disordered domains. As known from the previous studies using fluorescently labelled PEGylated cholesterol, DOPE and DSPE^5,17,18^, saturated DSPE prefers ordered domains, unsaturated DOPE prefers disordered domains, as expected from their native counterparts and cholesterol partitions into both phases with a slight ordered phase preference^17^. Directly labelled fluorescent DSPE without the PEG linker prefers disordered domains, highlighting the importance of PEG linker to preserve the native partitioning behaviour^5^. We first confirmed the partitioning of DOPE-PEG-ASR and DSPE-PEG-ASR in GUVs where DOPE-PEG-ASR prefers disordered domains (Lo %= 13 ± 5) and DSPE-PEG-ASR prefers ordered domains (Lo %= 67 ± 4, Figure 3A, B). In GPMVs, as previously suggested, partitioning of all analogues were relatively more ordered domain preferring due to more complex and natural ordering of phases in GPMVs^2,15,19^ (Figure 3B, C). DOPE-PEG-ASR still preferred disordered domains, yet with significantly higher Lo % (39 ± 3) while DSPE-PEG-ASR still preferred ordered domains with a similar extend (Lo %= 63 ± 4) (Figure 3A-C).

**Figure 3.**
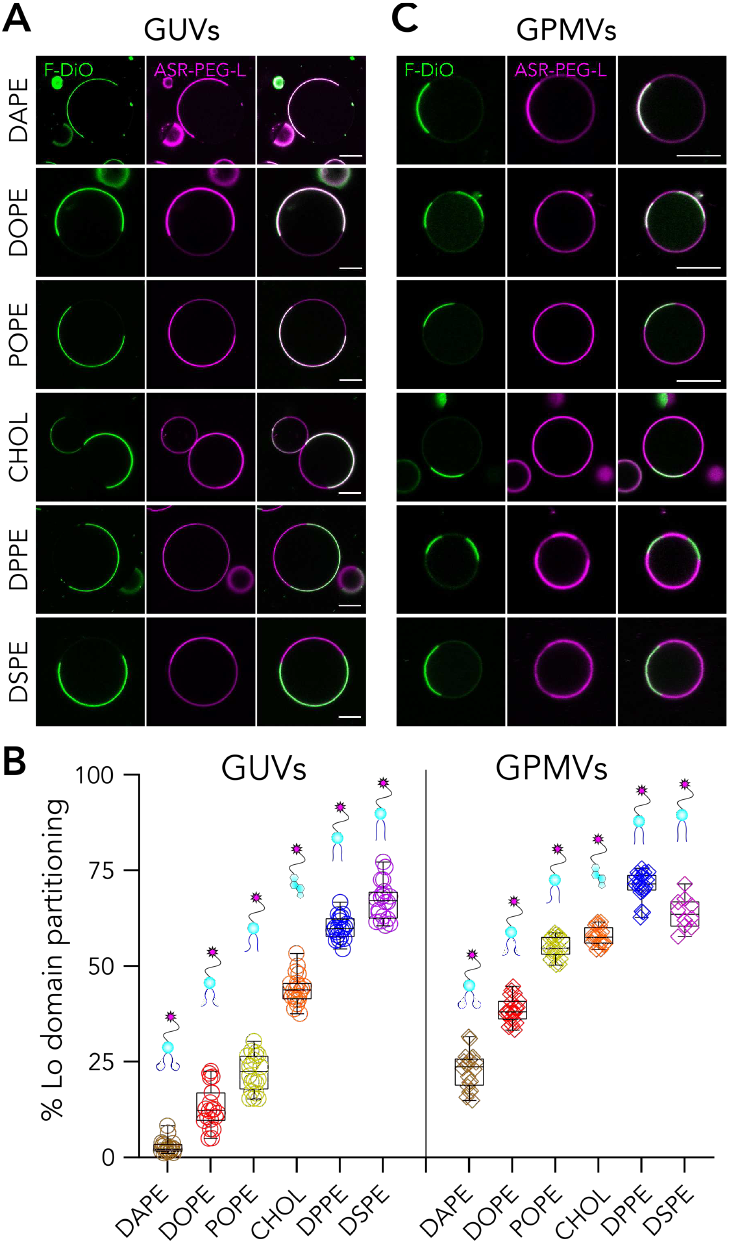
Partitioning of lipid analogues carrying PEG-2000 linker and labelled with Abberior Star Red (ASR) on the head group. Representative confocal images of A) phase separated GUVs and B) phase separated GPMVs doped with the fluorescence lipid analogues. C) Quantification of Lo % partitioning in GUVs and GPMVs. Each data point represents a vesicle. The graph is representative of three independent experiments. Scale bars are 10 µm.

It is common to use DOPE analogues as the disordered and the DSPE analogues as the ordered markers. However, having such a high Lo% in cell-derived GPMVs, DOPE seems suboptimal to use as the disordered marker in cellular membranes. To this end, we tested other lipid structures. DAPE is a polyunsaturated lipid (Figure 2A) and presumably a better Ld-phase lipid. When we labelled DAPE with the same strategy, the resulting analogue (DAPE-PEG-ASR) partitioned almost exclusively in disordered domains in GUVs (Lo %= 3 ± 2) and in GPMVs (Lo %= 22 ± 4). As an alternative to DSPE, we also tested DPPE which is a fully saturated lipid with shorter acyl chain length than DSPE. Surprisingly, compared to DSPE-PEG-ASR, DPPE-PEG-ASR partitioned less into the ordered domains in GUVs (Lo %= 60 ± 3) but more into the ordered domains in cell-derived GPMVs (Lo %= 72 ± 3). This suggests that DPPE might be a more suitable probe as an ordered phase marker in complex cellular membrane environment.

Interestingly, POPE has a hybrid acyl chain; a 16-carbon saturated chain and an 18-carbon mono-unsaturated chain. As shown above, fully saturated sixteen carbon DPPE-PEG-ASR prefers ordered domains while mono-unsaturated 18-carbon DOPE-PEG-ASR prefers disordered domains. Since POPE is a mixture of these two, it is not trivial to predict its phase behaviour, especially in complex live cell environment. Previous studies in fully synthetic membranes often used PO lipids as the disordered phase lipids^20^. This is mostly because when PO lipid (e.g., POPC) is mixed with a saturated lipid (e.g., DPPC or sphingomyelin) and cholesterol, phase separated membranes are formed where the disordered domains are thought to be mainly comprised of POPC^20^. However, we need to re-evaluate the partitioning of such lipids in more complex settings, such as in GPMVs. To this end, we synthesized POPE-PEG-ASR and tested its partitioning in GUVs and GPMVs. In GUVs, it partitioned into the Ld phase (Lo %= 22 ± 5) as expected. However, quite surprisingly, it partitioned slightly into the Lo phase in GPMVs (Lo %= 55 ± 3). This unexpected observation suggests a revision on the definition of an ordered vs. disordered lipids in complex cellular membranes.

In summary, our data suggest a correlation between the partitioning behaviour of probes with their internalization: more disordered preferring lipids get internalized more in cells except for DSPE (which potentially stems from the acyl chain length mismatch that will be discussed later).

### Time lapse and super-resolution imaging with DPPE-PEG-ASR

Having identified DPPE-PEG-ASR as an ideal probe to label the plasma membrane, we used it for the advanced applications such as time lapse imaging and super-resolution imaging. For time-lapse imaging, probes need to be photo-stable with minimal photobleaching. To test the suitability of these probes for time lapse microscopy, we imaged the stained U-2 OS cells continuously for 50 frames and compared the bleaching with a widely used plasma membrane dye, CellMask Deep Red. While the initial signal for both probes were similar, CellMask bleached significantly more than DPPE-PEG-ASR during 50-frame imaging (Figure 4A, B).

**Figure 4.**
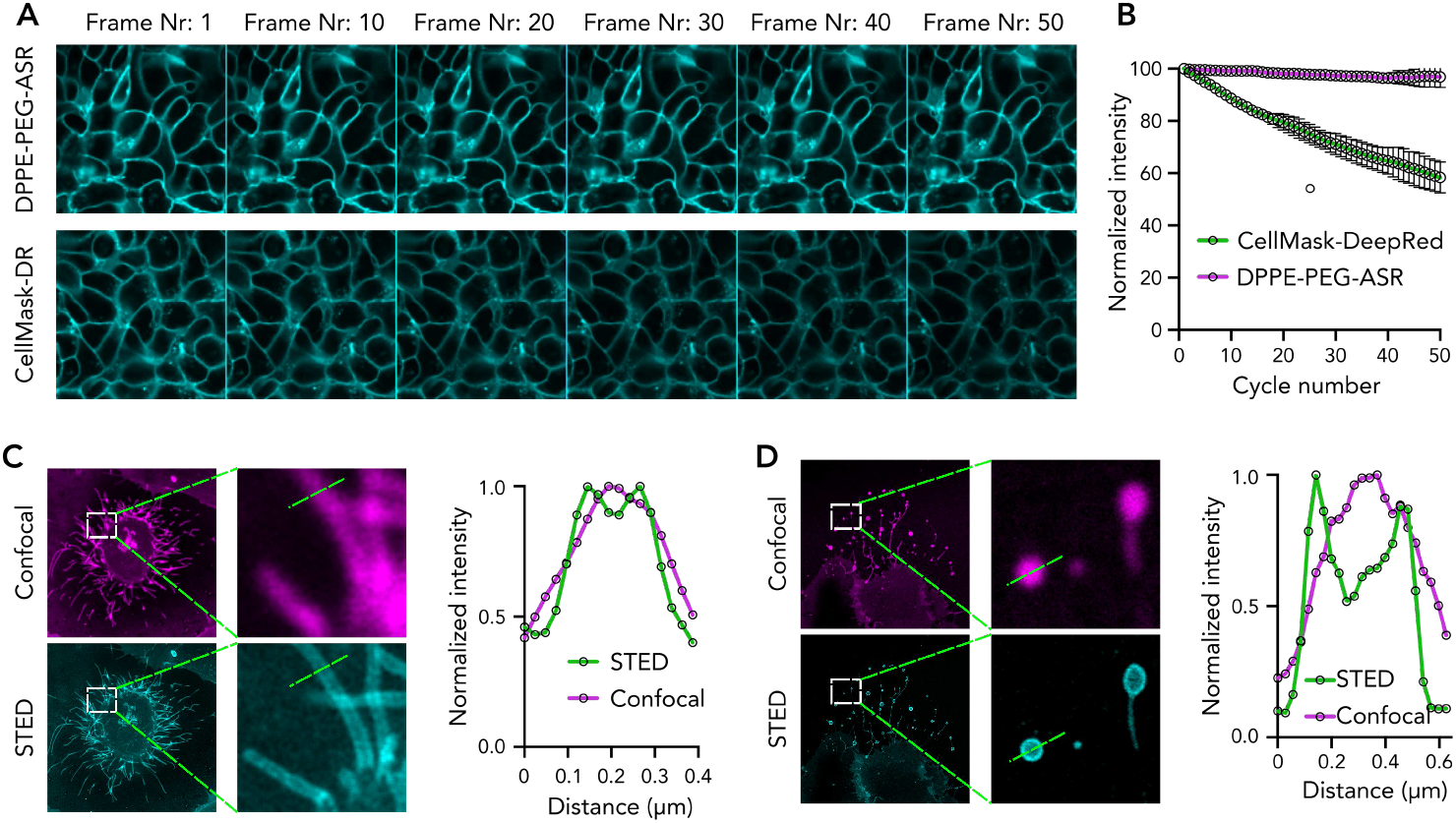
Advanced imaging with directly labelled lipid analogues. A) Timelapse imaging of DPPE-PEG-ASR and CellMask Deep Red with confocal microscopy. B) Quantification of photobleaching. C, D) STED images of U-2 OS cells labelled with DPPE-PEG-ASR and intensity profile of the lines shown in the image.

ASR has extensively been used for super-resolution stimulated emission depletion (STED) imaging^5,21^. To test whether our new probes can be used for super-resolution STED imaging of membranes, we performed STED imaging on U-2 OS cells stained with DPPE-PEG-ASR. To show the resolution enhancement, we imaged nanoscale membrane structures such as membrane tubes (Figure 4C) and migrasomes (Figure 4D). In both cases, STED combined with DPPE-PEG-ASR resolved the nanoscale structures as shown in the intensity profiles (Figure 4C, D).

This data shows that directly labelled DPPE-PEG-ASR is a promising alternative to available membrane probes for advanced membrane imaging. It also suggests that DPPE is a better candidate than other lipids we tested as the base lipid to be used for membrane labelling probes. However, this statement should be supported by evaluating other labelling approaches. For this reason, in the next section, we will study Azide-functionalized lipids.

### Membrane incorporation efficiency and partitioning of lipids labelled with Azide-DBCO conjugation

While the direct fluorescence labelling is convenient to label lipids, it still has some disadvantages. For example, it is not modular: to change the fluorophore, a new molecule has to be synthesized with the new fluorophore which is expensive and time-consuming. Moreover, when cells are incubated with the directly labelled fluorescent lipid analogues, their behaviour (e.g., localization, trafficking) is affected both by the lipid and the fluorescent tag. An alternative approach is functionalizing the lipid as minimally as possible with a small chemical moiety, incubate the cells with this minimally modified lipid analogue and only after the incorporation attach the fluorophore via click chemistry. Click chemistry has become an efficient way to generate biomolecular conjugates. A common click chemistry strategy for labelling lipids is Azide-DBCO conjugation. This approach is now extensively used for membrane labelling and functionalisation such as in the targeted lipid nanoparticles ^22,23^. Therefore, it is again crucial to test whether different lipid structures will be better suited to use for better membrane incorporation and labelling.

To apply this strategy for membrane labelling, azide functionalized lipids can be incorporated into the membranes followed by fluorescent DBCO addition. Alternatively, azide lipids and DBCO can be pre-mixed, and the resulting complex can be added to the membranes. Azide-DBCO labelling has the advantage of flexible fluorophore selection compared to direct fluorescence labelling. DBCO is available with several fluorescent tags, therefore one can choose the most appropriate dye for the planned experiments (e.g., super-resolution dyes, photo-switchable dyes etc.) without the need of resynthesizing the lipid analogue. Moreover, Azide-DBCO coupling can be used to attach other macromolecules to the membrane, such as antibodies, by functionalising them with either DBCO or Azide.

To test the suitability of such lipids for membrane labelling, we used azide-functionalized DOPE, DPPE and DSPE analogues, all carrying a 2K-PEG in between (Figure 1B). The selection of lipids was motivated by two intriguing observations discussed above both of which need confirmation with different labelling strategies. First, it was surprising to see DOPE-PEG-ASR exhibiting ≈40% Lo partitioning in GPMVs. Second intriguing observation was higher ordered domain partitioning of DPPE-PEG-ASR than DSPE-PEG-ASR which has longer saturated acyl chains. To evaluate the partitioning of Azide-functionalized lipids in GUVs and GPMVs, we pre-clicked Azide-functionalized lipids with DBCO with the same molar amount in a tube and later incubated the complex with GUVs or GPMVs. First, we observed a higher level of background compared to direct fluorescence labelling, which is presumably due to both unbound DBCO and un-incorporated lipid-Azide-DBCO complex (Figure 5A). Moreover, to be able to achieve the same level of signal as in the directly labelled lipids, we had to use substantially more lipids presumably owing to suboptimal clicking and incorporation efficiency. Interestingly, incorporation efficiency varied for different lipids. Both in GUVs and in GPMVs, the most efficiently incorporated lipid was DPPE-PEG-Azide-DBCO-Alexa647, followed by DOPE-PEG-Azide-DBCO-Alexa647 (Figure 5B). DSPE-PEG-Azide-DBCO647 incorporated poorly compared to the other analogues (Figure 5A).

**Figure 5.**
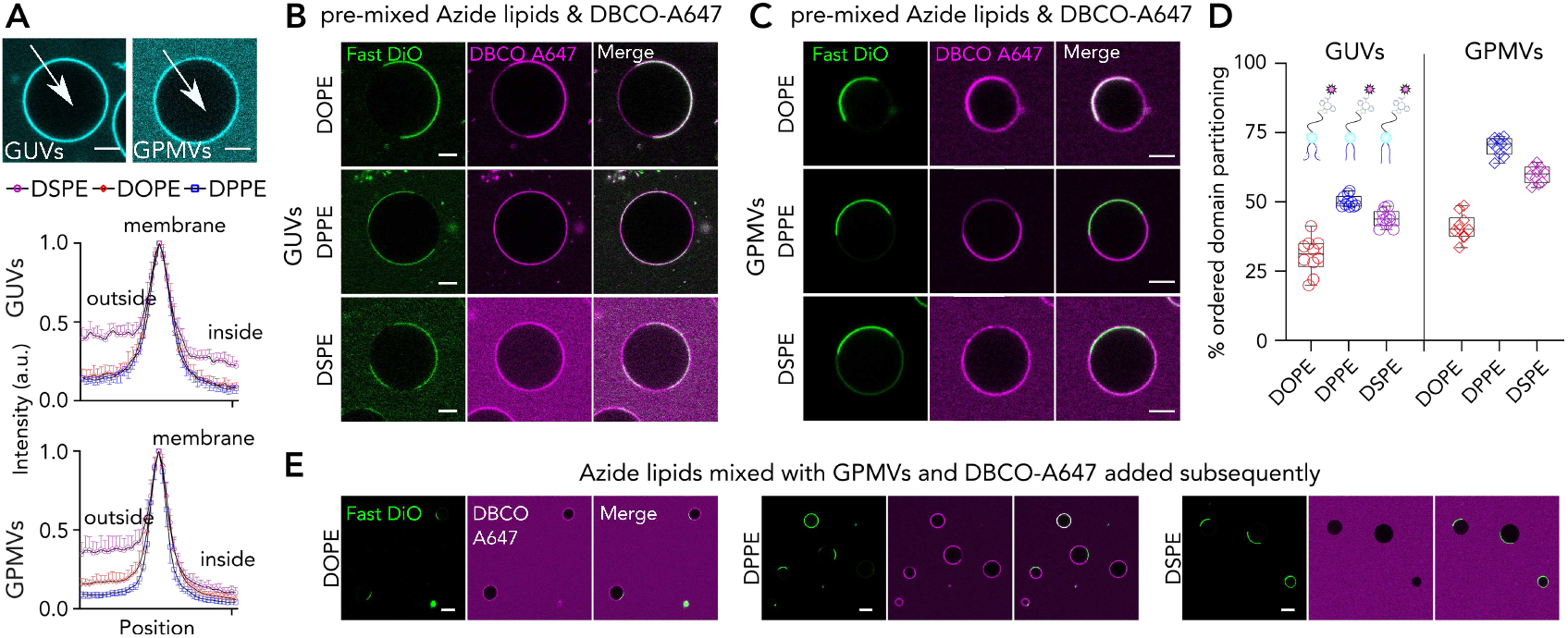
Membrane incorporation and partitioning of lipid analogues carrying PEG-Azide on the headgroup and labelled with fluorescent DBCO. A) Incorporation efficiency of lipid analogues in GUVs and GPMVs measured by the internal and external signal compared to the membrane signal. Large outside signal suggest less efficient incorporation. Data is the mean of ten vesicles with error bars representing the standard deviation. Representative confocal images of phase separated B) GUVs and C) GPMVs doped with the lipid analogues. D) Quantification of Lo % partitioning in GUVs and GPMVs. Each data point represents a vesicle. The graph is representative of three independent experiments. E) Representative confocal images of GPMVs doped with azide-functionalized lipids first, and DBCO-647 treated later. Scale bars are 10 µm.

DOPE-PEG-Azide-DBCO-Alexa647, similar to DOPE-PEG-ASR, partitioned into Ld with Lo % of 30 ± 6 in GUVs and 41 ± 5 in GPMVs. This further strengthens that DOPE is not optimal to be used as the disordered membrane marker in complex membranes. The poor incorporation of DSPE-PEG-Azide-DBCO-Alexa647 presumably stems from the hydrophobic mismatch of long acyl chain lipids. In such a case, the partitioning would also be affected as the lipid is not properly inserted into the membrane. Indeed, we observed that DSPE-PEG-Azide-DBCO-Alexa647 exhibited lower Lo partitioning compared to DPPE-PEG-Azide-DBCO-Alexa647 both in GUVs and in GPMVs (Figure 5B-D)as we observed for the DSPE-PEG-ASR. To test how pre-clicking changes the labelling efficiency, we added only the azide-functionalized lipids on the membranes and subsequently added the DBCO. The inefficient incorporation of DSPE-PEG-Azide was even more pronounced when we first incorporated the lipids into GPMVs and then added DBCO-Alexa647 subsequently (Figure 5E).

Overall, this labelling strategy has the advantage of flexible fluorophore choice, but for model membrane studies, where sample washing is technically challenging, has the major disadvantage of background from unbound molecules. For cellular experiments, however, this labelling strategy is a viable approach. DPPE again showed better performance as the base lipid compared to the other we tested.

### Imaging live, fixed and permeabilized cells using Azide functionalized lipids

For live cell labelling, we pre-clicked the Azide lipids with DBCO in a test tube and incubated the complex with cells. After incubation, the cells were washed to get rid of the unbound molecules. While all probes were observed in the plasma membrane of both Jurkat and U-2 OS cells (Figure 6A, B), incorporation efficiency varied depending on the lipid structure. DPPE-PEG-Azide-DBCO-Alexa647 showed the most efficient staining followed by DOPE-PEG-Azide-DBCO-Alexa647. DSPE-PEG-Azide-DBCO-Alexa647 showed the poorest incorporation among the probes (Figure 6A-C). We also incubated the cells with the analogues for 3h to test the internalization (Figure 3B, lower panels; intensity adjusted to have visually similar intensities between probes). While it is technically challenging to quantify the internalization due to initial difference in incorporation, visually DPPE-PEG-Azide-DBCO-Alexa647 exhibits the lowest internalization as seen from the lack of puncta inside the cells unlike DSPE and DOPE analogues. Finally, we showed that DPPE-PEG-Azide-DBCO-Alexa647 remains in the plasma membrane post-fixation (Figure 6D).

**Figure 6.**
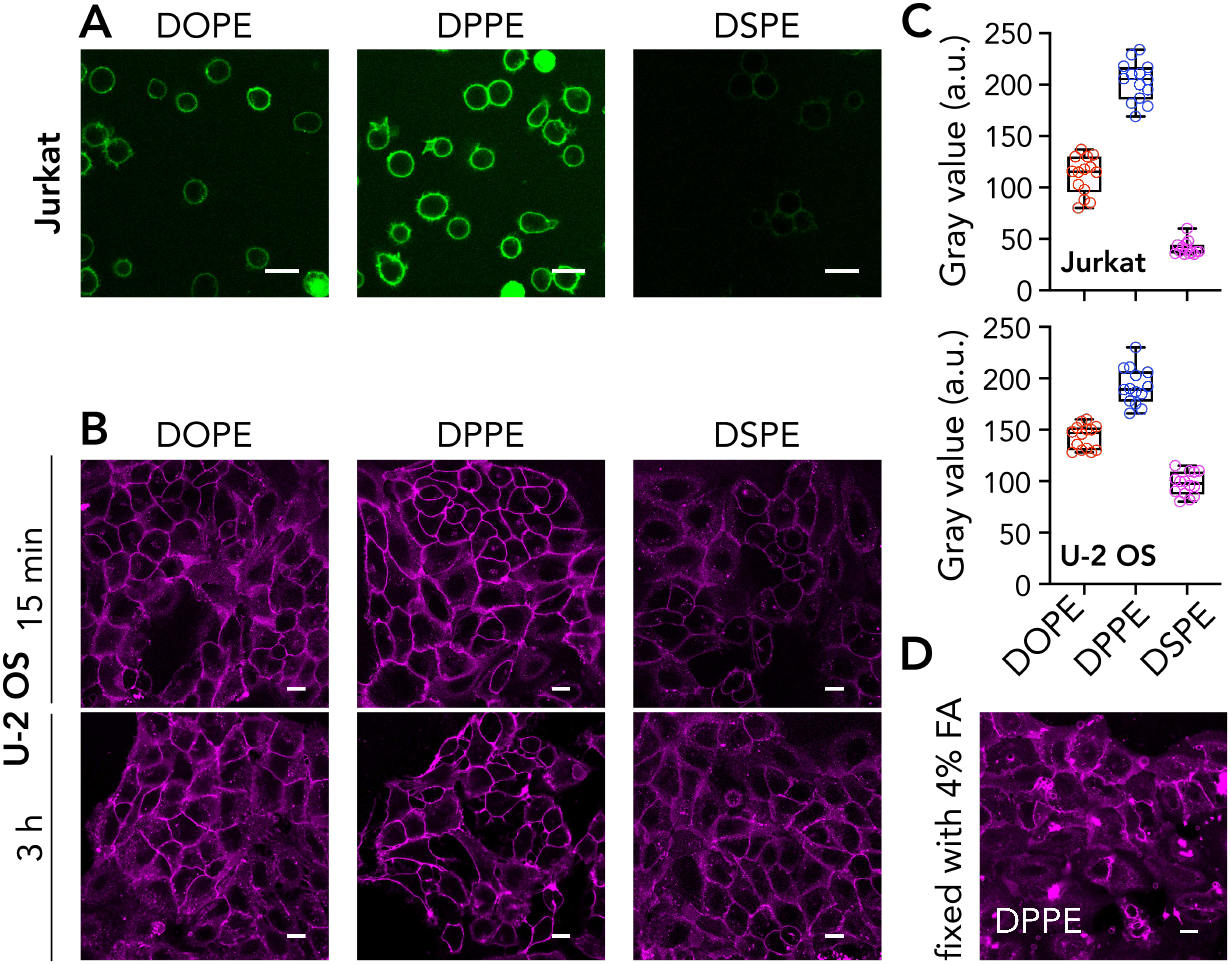
Imaging live, fixed and permeabilized cells using Azide functionalized lipids. A) Jurkat cells incubated with Azide-functionalized DOPE-PEG, DPPE-PEG and DSPE-PEG and stained with DBCO-AF647. B) U-2 OS cells incubated with Azide-functionalized DOPE-PEG, DPPE-PEG and DSPE-PEG and stained with DBCO-AF647; top panel for 15 minutes after incubation, bottom panel for 3h after incubation. Bottom panels are intensity-adjusted to make internalization visually clear. C) Quantification of labelling efficiency. D) Fixed U-2 OS cells stained with DPPE-PEG-Azide-DBCO-AF647.

## Conclusions

Here, we evaluated different lipid analogues to label the plasma membrane effectively for live, fixed and permeabilized cells. We tested structurally different lipids and two different labelling approaches and identified DPPE as the most suitable lipid to be used for labelling the plasma membrane of mammalian cells. Both directly labelled DPPE-PEG-ASR and functionalized DPPE-PEG-Azide probes labelled the plasma membrane more efficiently than the other probes we tested. Moreover, DPPE remained in the plasma membrane longer for time-lapse experiments. This is presumably due to the ordered domain partitioning of DPPE since disordered membrane probes labelled the same approach showed more internalisation. Interestingly, DSPE, which has two saturated acyl chains with 18 carbons partitioned less into the ordered domains in cell-derived GPMVs, suggesting a mismatch in acyl chain length. In line with this, DSPE also internalized more compared to DPPE. Therefore, DPPE analogues can be used to study plasma membrane structure and dynamics, as we demonstrated for time lapse and super resolution imaging. For direct labelling, we used Abberior Star Red since it is a photostable probe, hence its bleaching is minimal allowing for advanced imaging. Similarly, DPPE-PEG-Azide probe can be clicked to a photostable dye for advanced imaging.

Here, we showed the application of the probes in eukaryotic cells and model membranes, but these data will also help in designing better probes for other structures such as lipid nanoparticles and extracellular vesicles. Additionally, since we showed lipid-dependent internalization, these probes can also be used to study endocytosis of different molecules to test whether their endocytosis is dependent on the lipid environment. We also presented new previously untested probes with a spectrum of partitioning ranging from a perfectly disordered lipid analogue (DAPE) to a more equipartitioning analogue (POPE) to an ordered domain partitioning analogue (DPPE). This will also be important for the membrane biology field where membrane domains and lipid dependent membrane dynamics are studied.

Overall, our systematic evaluation will be a useful guideline for the membrane biology field with these new tools to study membrane structure and dynamics. Moreover, the tools we present here will be crucial for the spatial biology field where membrane labelling has always been challenging.

## Data Availability

All data will be available upon publication in Figshare.

## Author contributions

ES designed and performed the experiments, analysed the data and wrote the manuscript.

## Acknowledgements

I would like to thank the member of CSI:Nano Lab and ALM SciLifeLab team for discussions. ES has been supported by Swedish Research Council Grants (grant no. 2020-02682, 2024-02993 and 2024-00289), Wellcome Leap’s Dynamic Resilience Program (jointly funded by Temasek Trust), Karolinska Institutet (2024-03250; 2024-03341; 2022-00803; 2020-00997), Cancer Research KI (2024-03488), Human Frontier Science Program (RGP0025/2022), Longevity Impetus Grant from Norn Group, Hevolution Foundation and Rosenkranz Foundation.

## References

1 A. S. Klymchenko and R. Kreder, Chemistry & Biology, 2014, 21, 97–113.

2 E. Sezgin, I. Levental, M. Grzybek, G. Schwarzmann, V. Mueller, A. Honigmann, V. N. Belov, C. Eggeling, U. Coskun, K. Simons and P. Schwille, Biochimica Et Biophysica Acta-Biomembranes, 2012, 1818, 1777–1784.

3 S. Rissanen, M. Grzybek, A. Orlowski, T. Rog, O. Cramariuc, I. Levental, C. Eggeling, E. Sezgin and I. Vattulainen, Frontiers in physiology, 2017, 8, 252.

4 P. Sengupta, A. Hammond, D. Holowka and B. Baird, Biochimica Et Biophysica Acta-Biomembranes, 2008, 1778, 20–32.

5 A. Honigmann, V. Mueller, S. W. Hell and C. Eggeling, Faraday Discussions, 2013, 161, 77–89.

6 A. B. Neef and C. Schultz, Angewandte Chemie International Edition, 2009, 48, 1498–1500.

7 A. L. L. Matos, F. Keller, T. Wegner, C. E. C. del Castillo, D. Grill, S. Kudruk, A. Spang, F. Glorius, A. Heuer and V. Gerke, Commun Biol, 2021, 4, 1–11.

8 R. Blankenburg, P. Meller, H. Ringsdorf and C. Salesse, Biochemistry, 1989, 28, 8214–8221.

9 S. Henry, E. Williams, K. Barr, E. Korchagina, A. Tuzikov, N. Ilyushina, S. A. Abayzeed, K. F. Webb and N. Bovin, Sci Rep, 2018, 8, 2845.

10 N. O. Fischer, C. D. Blanchette, B. A. Chromy, E. A. Kuhn, B. W. Segelke, M. Corzett, G. Bench, P. W. Mason and P. D. Hoeprich, Bioconjugate Chem., 2009, 20, 460–465.

11 G. G. Chikh, W. M. Li, M.-P. Schutze-Redelmeier, J.-C. Meunier and M. B. Bally, Biochimica et Biophysica Acta (BBA) - Biomembranes, 2002, 1567, 204–212.

12 W. Zheng, M. Schürz, R. J. Wiklander, O. Gustafsson, D. Gupta, R. Slovak, A. Traista, A. Coluzzi, S. Roudi, A. Barone, D. Farcas, E. Kyriakopoulou, V. Galli, H. Sharma, N. Meisner-Kober, M. Honcharenko and S. E. L. Andaloussi, J Control Release, 2023, 357, 630–640.

13 A. J. Garcia-Saez, D. C. Carrer and P. D.-:10.1007/978-1-60761-447-0_33 Schwille, Liposomes: Methods and Protocols, Vol 2:BIOLOGICAL MEMBRANE MODELS, 2010, 493–508.

14 E. Sezgin, H.-J. Kaiser, T. Baumgart, P. Schwille, K. Simons and I. Levental, Nature Protocols, 2012, 7, 1042–1051.

15 E. Sezgin, T. Gutmann, T. Buhl, R. Dirkx, M. Grzybek, U. Coskun, M. Solimena, K. Simons, I. Levental and P. Schwille, PloS one, 2015, 10, e0123930.

16 A. El-Sayed and H. Harashima, Mol Ther, 2013, 21, 1118–1130.

17 A. Honigmann, V. Mueller, H. Ta, A. Schoenle, E. Sezgin, S. W. Hell and C. Eggeling, Nature Communications, 2014, 5, 5412–5412.

18 E. Mobarak, M. Javanainen, W. Kulig, A. Honigmann, E. Sezgin, N. Aho, C. Eggeling, T. Rog and I. Vattulainen, Biochimica et biophysica acta, DOI:10.1016/j.bbamem.2018.07.003.

19 H. J. Kaiser, D. Lingwood, I. Levental, J. L. Sampaio, L. Kalvodova, L. Rajendran and K. Simons, Proceedings of the National Academy of Sciences of the United States of America, 2009, 106, 16645–16650.

20 S. L. Veatch and S. L. Keller, Physical Review Letters, DOI:10.1103/PhysRevLett.94.148101.

21 A. Barbotin, I. Urbancic, S. Galiani, C. Eggeling, M. Booth and E. Sezgin, Biophys J, DOI:10.1016/j.bpj.2020.04.006.

22 Y. Sakurai, N. Abe, K. Yoshikawa, R. Oyama, S. Ogasawara, T. Murata, Y. Nakai, K. Tange, H. Tanaka and H. Akita, Journal of Controlled Release, 2022, 349, 379–387.

23 X. Li, S. Weller, G. Clergeaud and T. L. Andresen, Biotechnol J, 2024, 19, e2300339.

